# A complex multi-locus, multi-allelic genetic architecture underlying the long-term selection-response in the Virginia body weight line of chickens

**DOI:** 10.1101/098160

**Authors:** Yanjun Zan, Zheya Sheng, Lars Rönnegård, Christa F. Honaker, Paul B. Siegel, Örjan Carlborg

**Author notes:** Author emails: YZ ZS LR, CFH PBS ÖC. These authors contributed equally (Yanjun Zan, Zheya Sheng).

## Abstract

The ability of a population to adapt to changes in their living conditions, whether in nature or captivity, often depends on polymorphisms in multiple genes across the genome. In-depth studies of such polygenic adaptations are difficult in natural populations, but can be approached using the resources provided by artificial selection experiments. Here, we dissect the genetic mechanisms involved in long-term selection responses of the Virginia chicken lines, populations that after 40 generations of divergent selection for 56-day body weight display a nine-fold difference in the selected trait. In the F15 generation of an intercross between the divergent lines, 20 loci explained more than 60% of the additive genetic variance for the selected trait. We focused particularly on seven major QTL and found that only two fine-mapped to single, bi-allelic loci; the other five contained linked loci, multiple alleles or were epistatic. This detailed dissection of the polygenic adaptations in the Virginia lines provides a deeper understanding of genome-wide mechanisms involved in the long-term selection responses. The results illustrate that long-term selection responses, even from populations with a limited genetic diversity, can be polygenic and influenced by a range of genetic mechanisms.

## Introduction

Experimentally adapted populations are powerful resources for dissecting the genetic basis of adaptation (Hill 2005). These closed populations, subjected to long-term artificial selection for clearly defined adaptive traits, will accumulate adaptive genetic variations at a more rapid rate than natural populations. By mapping individual loci contributing to selection response in these populations, it is possible to gain fundamental insights on the genetic architecture of complex traits and their contributions to adaptation and evolution. As highlighted in Churchill (Churchill 2016), efforts to map quantitative trait loci (QTL) have often revealed an almost overwhelming complexity. Fine-mapping efforts have been challenged by the highly polygenic architecture of traits, where each of the contributing loci often encompass additional complexities, including multiple tightly linked variants with small or large, additive or epistatic genetic effects (; Brandt et al. 2016; Holland 2007; Laurie et al. 2004; Mott and Flint 2013). Therefore, there is a shortage of studies that provide a deeper understanding about the genetic architectures contributing to the adaptive response to longterm selection (Burke 2012).

Modern genomics allows cost-efficient, in-depth characterization of genetic variation across many loci, or entire genomes, in large populations. This provides opportunities to design studies that make efficient use of selected populations to gain a genome-wide perspective on the genetic architecture of long-term selection response (Burke 2012; Johansson et al. 2010; Pettersson et al. 2013) or to dissect the complexity within and across the loci that contribute to adaptations for highly polygenic traits (Brandt et al. 2016; Chan et al. 2012; Sheng et al. 2015). Here, we describe the simultaneous fine-mapping of nine QTL contributing to longterm selection responses in the Virginia chicken lines. These lines have been under bidirectional selection for 56-day body weight (Dunnington and Siegel 1996; Dunnington et al. 2013; Márquez et al. 2010). After 40 generations of selection, the High- (HWS) and Low- (LWS) weight selected lines differed 9-fold for the selected trait. In the current generation (S59) the lines differ by more than 16-fold with many loci contributing to this difference (Besnier et al. 2011; Carlborg et al. 2006; Jacobsson et al. 2005; Johansson et al. 2010; Pettersson et al. 2011; Sheng et al. 2015; Wahlberg et al. 2009) with likely contributions by allelic heterogeneity (Besnier *et al.* 2011; Brandt *et al.* 2016) and epistasis (Pettersson et al. 2011).

Reported here is our most recent progress in dissection of the genetic architecture underlying the extreme adaptations in the Virginia body weight chicken lines. Seven major QTL, mapped and confirmed in earlier reports (Besnier et al. 2011; Brandt et al. 2016; Jacobsson et al. 2005; Wahlberg et al. 2009), were fine-mapped using data from the F_15_ generation of an Advanced Intercross Line bred from HWS and LWS founders from generation 40. We performed a multi-locus, association-based fine-mapping analysis, and explored the changes in allele frequencies across the QTL during ongoing selection in the parental lines. Revealed was the genetic basis of several QTL where earlier statistical analyses suggested more complex underlying genetic architectures than a single causal bi-allelic locus (Besnier et al. 2011; Brandt et al. 2016). In particular, examples are provided of how tight linkage, multiple alleles from the founder-lines, and epistatic interactions between fine-mapped loci contribute to the QTL effects. In total, seven QTL were fine-mapped into 10 contributing loci, and another 10 loci earlier reported elsewhere in the genome were confirmed (Sheng et al. 2015). Together, these 20 loci explain more than 60% of the additive genetic variance in the population, illustrating how this study provides one of the more comprehensive dissections of a complex adaptive trait in animals. By developing and using new statistical approaches to analyse this novel resource population, we provide insights on the genetic mechanisms contributing to long-term responses to selection that could guide future work to further understanding of the genetic basis of adaptation and evolution.

## Results

### Seven of nine QTL were replicated in the F_15_ generation of the Advanced Intercross Line

Our fine-mapping analysis of nine QTL affecting 56-day body weight in the Virginia chicken lines (Besnier et al. 2011; Jacobsson et al. 2005; Wahlberg et al. 2009;) used genotypic and phenotypic data from the F_15_ generation of an Advanced Intercross between HWS and LWS founders from generation 40. A multi-locus backward-elimination analysis with bootstrapping to correct for population structure allowed scanning for independent associations across 216 markers in the nine QTL, and 218 markers from selective-sweeps located elsewhere in the genome (Sheng et al. 2015). In total, 24 SNP markers with statistically independent associations to 56-day body-weight were identified at 20% False Discovery Rate (FDR). Of these, 13 were located in seven of the nine QTL (Table 1), and 11 in selective-sweeps elsewhere in the genome (Supplemental Table S1).

**Table 1.**
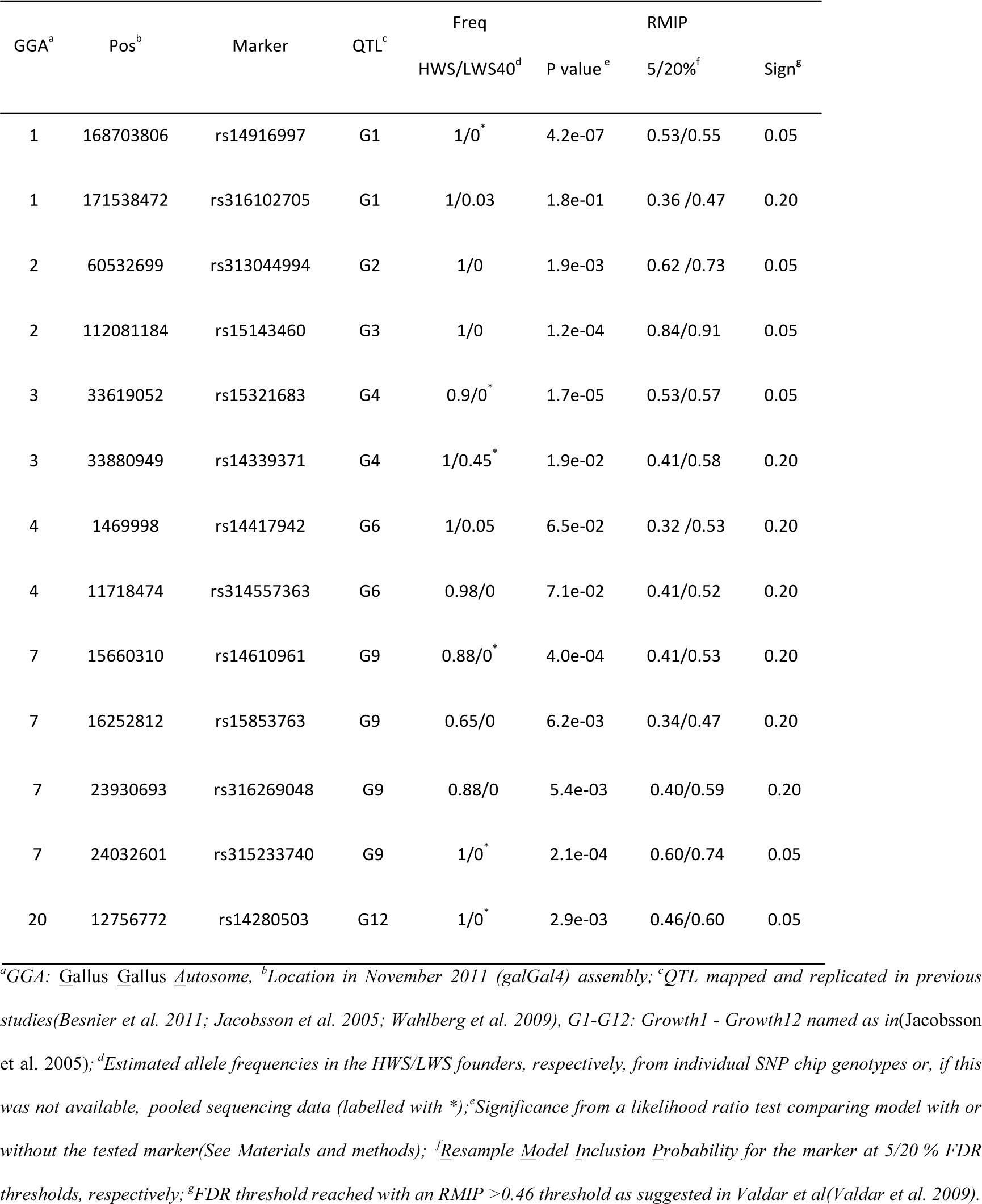
In the nine fine-mapped QTL, 13 SNP markers are associated with 56-day body-weight in generation F_15_ of the Virginia Advanced Intercross Line. Outside of these QTL, associations to 11 additional markers are detected (Supplemental Table S1).

| GGA<sup>a</sup> | Pos<sup>b</sup> | Marker | QTL<sup>c</sup> | Freq |  | RMIP |  |
| --- | --- | --- | --- | --- | --- | --- | --- |
|  |  |  |  | HWS/LWS40<sup>d</sup> | P value<sup>e</sup> | 5/20%<sup>f</sup> | Sign<sup>g</sup> |
| 1 | 168703806 | rs14916997 | G1 | 1/0<sup>*</sup> | 4.2e-07 | 0.53/0.55 | 0.05 |
| 1 | 171538472 | rs316102705 | G1 | 1/0.03 | 1.8e-01 | 0.36 /0.47 | 0.20 |
| 2 | 60532699 | rs313044994 | G2 | 1/0 | 1.9e-03 | 0.62 /0.73 | 0.05 |
| 2 | 112081184 | rs15143460 | G3 | 1/0 | 1.2e-04 | 0.84/0.91 | 0.05 |
| 3 | 33619052 | rs15321683 | G4 | 0.9/0<sup>*</sup> | 1.7e-05 | 0.53/0.57 | 0.05 |
| 3 | 33880949 | rs14339371 | G4 | 1/0.45<sup>*</sup> | 1.9e-02 | 0.41/0.58 | 0.20 |
| 4 | 1469998 | rs14417942 | G6 | 1/0.05 | 6.5e-02 | 0.32 /0.53 | 0.20 |
| 4 | 11718474 | rs314557363 | G6 | 0.98/0 | 7.1e-02 | 0.41/0.52 | 0.20 |
| 7 | 15660310 | rs14610961 | G9 | 0.88/0<sup>*</sup> | 4.0e-04 | 0.41/0.53 | 0.20 |
| 7 | 16252812 | rs15853763 | G9 | 0.65/0 | 6.2e-03 | 0.34/0.47 | 0.20 |
| 7 | 23930693 | rs316269048 | G9 | 0.88/0 | 5.4e-03 | 0.40/0.59 | 0.20 |
| 7 | 24032601 | rs315233740 | G9 | 1/0* | 2.1e-04 | 0.60/0.74 | 0.05 |
| 20 | 12756772 | rs14280503 | G12 | 1/0* | 2.9e-03 | 0.46/0.60 | 0.05 |

<sup>a</sup>GGA: *Gallus Gallus Autosome*, <sup>b</sup>Location in November 2011 (*galGal4*) assembly; <sup>c</sup>QTL mapped and replicated in previous studies (Besnier et al. 2011; Jacobsson et al. 2005; Wahlberg et al. 2009), *G1-G12: Growth1 - Growth12* named as in (Jacobsson et al. 2005); <sup>d</sup>Estimated allele frequencies in the HWS/LWS founders, respectively, from individual SNP chip genotypes or, if this was not available, pooled sequencing data (labelled with \*); <sup>e</sup>Significance from a likelihood ratio test comparing model with or without the tested marker (See Materials and methods); <sup>f</sup>Resample Model Inclusion Probability for the marker at 5/20 % FDR thresholds, respectively; <sup>g</sup>FDR threshold reached with an RMIP >0.46 threshold as suggested in Valdar et al (Valdar et al. 2009).

### Fine-mapping of the seven replicated QTL

We tested for associations to each of the SNP markers within each of the fine mapped QTL in the AIL F_15_ generation. This analysis was performed to obtain QTL profiles in this generation for comparison with earlier results from the F_2_ generation (Wahlberg et al. 2009). In this analysis, all markers detected at 20% FDR in multi-locus bootstrapping analysis were fitted as fixed effects to account for the multi-locus genetic architecture of 56-day body-weight in this population. The QTL profiles revealed that two QTL, *Growth2* on GGA2 and *Growth3* on GGA2; QTL names as in (Jacobsson et al. 2005), could be fine-mapped to single, bi-allelic loci (Figure 1; Table 1). More complex genetic architectures involving multiple linked loci, segregation of multiple alleles in at least one of the founder lines, or both were revealed in the other five QTL *(Growth1 on GGA1, Growth4 on GGA3, Growth6 on GGA4, Growth9 on GGA7* and *Growth12 on GGA20;* Table 1; Figure 2). Described in the sections below are more detailed results from the fine-mapping analyses.

**Figure 1.**
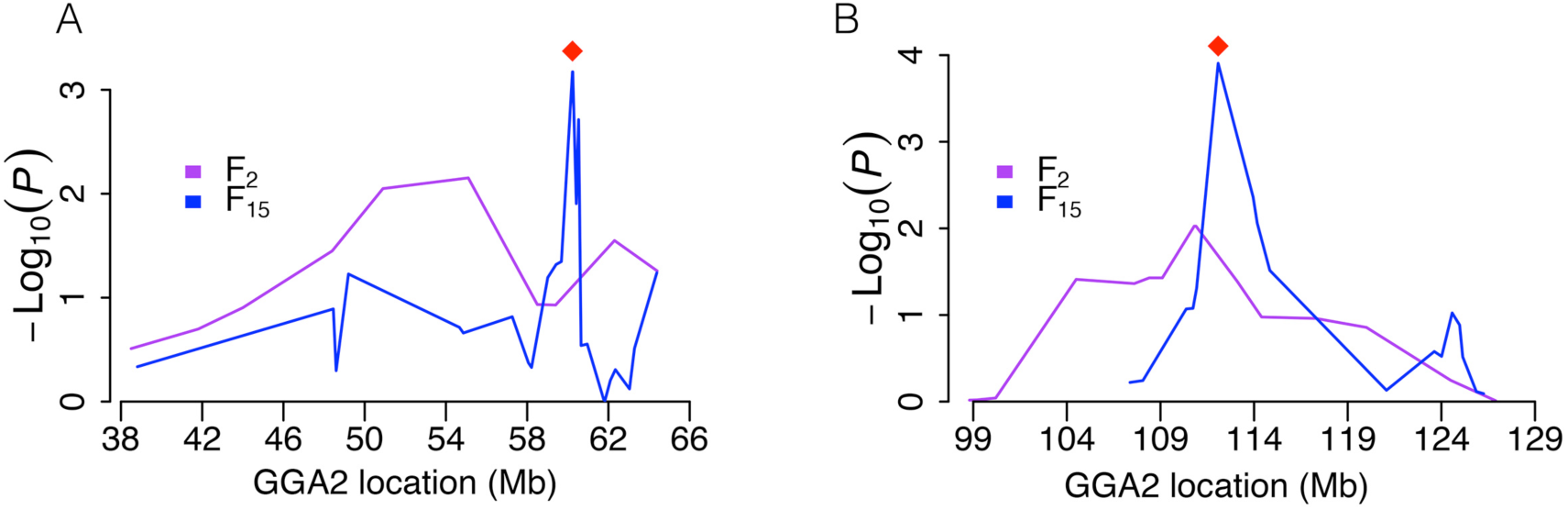
Statistical support curves from **QTL**-scans across the**QTL** *Growth2* **(A)** and *Growth3* **(B)** on GGA2. These QTL were originally detected in the F_2_ intercross generation (purple line) and fine-mapped here in the F_15_ generation (blue line) of an AIL. The lines connect the significances at the locations of genotyped markers in the two populations, from Wahlberg et al (Wahlberg et al. 2009) and this study, and the red diamonds highlight the marker retained in the multi-locus bootstrap-based backward-elimination analysis in the F_15_ generation (Table 1).

**Figure 2.**
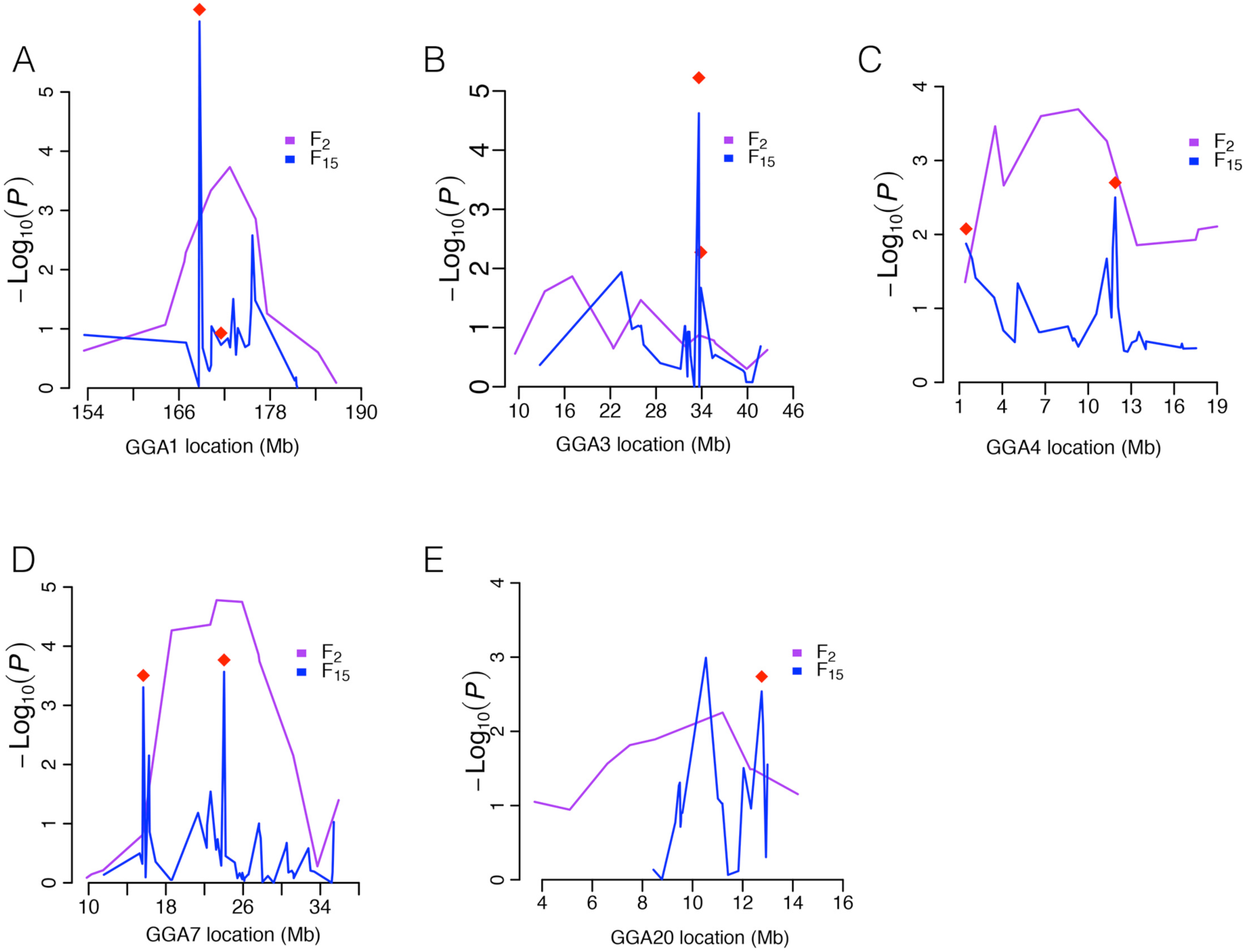
Statistical support curves from the QTL-scans across five QTL associated with 56-day body weight in the F_2_ intercross generation (purple) (Wahlberg et al. 2009) and fine mapped in the F_15_ generation of the AIL (blue; this study). The plots show **(A)** *Growth1* on GGA1, **(B)** *Growth4* on GGA3, **(C)** *Growth6* on GGA4, **(D)** *Growth9* on GGA7, and **(E)** Growth 12 on GGA20. The red diamonds indicate the significant markers in the bootstrap-based backward-elimination analysis in the F_15_ generation (Table 1).

### Two QTL fine-mapped to single locus with fixed alleles in the founder lines

The multilocus bootstrap-based backward-elimination analysis revealed only one 56-day body weight associated marker in each of the two QTL on GGA2 *(Growth2, Growth3;* Table 1). This result is consistent with that of the QTL-scan across *Growth2* that identifies a well-defined region with a peak at 60.5Mb (galGal4) to the marker retained in the backward-elimination analysis (Figure 1A). The fine-mapped association signal in the F_15_ is located ∼5 Mb away from, and slightly outside of, the QTL-peak identified in the original F_2_ analysis (Wahlberg et al. 2009) (59.4 Mb vs 55Mb in F_15_ and F_2_, respectively; Figure 1A). The additive genetic effects at the top associated markers in the F_2_ and F_15_ are, however, similar (less than 7g difference; Table 2).

**Table 2.** Estimates of the additive effects for the markers closest to the fine-mapped QTL-peaks in the F_2_ (Wahlberg et al. 2009) and F_15_ intercross generations (this study).

| Locus<sup>1</sup> | GGA<sup>2</sup> | QTL | Markers<sup>3</sup> | Position<sup>4</sup> | F<sub>2</sub><br>Additive<sup>5</sup> | F<sub>15</sub><br>Additive<sup>6</sup> |
| --- | --- | --- | --- | --- | --- | --- |
|  |  |  | (F<sub>15</sub>/F<sub>2</sub>) | (F<sub>15</sub>/F<sub>2</sub> Mb) | (a±SE) | (a±SE) |
| 1 | 1 | Growth1 | rs14916997/rs15502284 | 168.7/170.2 | 30.0± 8.5 | 26.2±5.8 |
| 2 | 1 | Growth1 | rs316102705/rs15502284 | 171.5/170.2 | 30.0± 8.5 | -6.9±5.9 |
| 3 | 2 | Growth2 | rs313044994/rs15105085 | 60.5/59.4 | 13.3± 8.5 | 20.9±7.1 |
| 4 | 2 | Growth3 | rs15143460/rs16107541 | 112.1/113.1 | 17.4± 8.5 | 21.4±6.3 |
| 5 | 3 | Growth4 | rs15321683/rs15321683 | 33.6/33.6 | 12.8± 8.6 | 21.1±5.6 |
|  |  |  | rs14339371/rs15321683 | 33.9/33.6 | 12.8± 8.6 | 9.9±6.5 |
| 6 | 4 | Growth6 | rs14417942/ADL0143 | 1.5/1.4 | 17.1± 8.5 | 10.1±6.0 |
| 7 | 4 | Growth6 | rs314557363/rs14428592 | 11.7/11.3 | 29.0± 8.4 | 10.6±6.1 |
| 8 | 7 | Growth9 | rs14610961/rs15852323 | 15.7/15.5 | 14.8±10.5 | 11.8±5.9 |
|  |  |  | rs15853763/rs15852323 | 16.3/15.5 | 14.8±10.5 | 0.8±5.7 |
| 9 | 7 | Growth9 | rs316269048/ADL0279 | 23.9/23.2 | 37.5± 8.6 | 1.8±5.6 |
|  |  |  | rs315233740/ADL0279 | 24.0/23.2 | 37.5± 8.6 | 12.7±5.5 |
| 10 | 20 | Growth12 | rs14280503/MC3R_PYRO | 12.8/12.4 | 18.1± 8.4 | 15.1±5.7 |

<sup>1</sup>Locus: Physically closed markers identified in the bootstrap-based backward-elimination analysis (<1 Mb apart) are clustered into a locus; <sup>2</sup>Gallus gallus Autosome; <sup>3</sup>Markers for which additive estimates are compared. In the F<sub>15</sub>, the markers were from the bootstrap-based, backward-elimination analysis and in the F<sub>2</sub> the marker in Wahlberg et al (Wahlberg et al. 2009) with the shortest physical distance (Mb galGal4) to the respective F<sub>15</sub> markers; <sup>4</sup>Location of markers in the Nov. 2011 (galGal4) assembly; <sup>5</sup>Additive genetic effect ± Standard Error estimated using a model including the genotype for the tested marker and the sex of the bird in the F<sub>2</sub> generation (Wahlberg et al. 2009); <sup>6</sup>Additive genetic effect ± Standard Error estimated in F<sub>15</sub> generation.

The QTL-scan across *Growth3* in the F_15_ generation identifies a single association peak at 112.1 Mb to the same marker detected in the multi-locus analysis (Figure 1B). The association signal is located approximately 2 Mb from the top associated marker in the F_2_ (Figure 1B). In *Growth3,* the peak marker in the F_2_ population was the closest one among all other peak markers to the fine-mapped location in the F_15_ generation, and the additive effects in the two generations differ only marginally (4g; Table 2). Thus, *Growth2* and *Growth3* QTL could be fine-mapped to narrow chromosomal regions using the data from the AIL F_15_ generation (Figure 1). The 1 LOD drop-off confidence intervals for *Growth2* and *Growth3* were 0.5 Mb (60.2-60.7) and 1.9 Mb (112.0-113.9), respectively.

### Two linked loci are revealed in four fine-mapped QTL

In four of the fine-mapped QTL*(Growthl on GGA1, Growth6 on GGA4, Growth9 on GGA7* and *Growth12 on GGA20;* Figure 2; Table 1), significant associations were detected to distant markers (> 2.5 Mb apart) in either the QTL-scan, the backward-elimination analysis or both in the F_15_ generation data. In addition to this, pairs of physically close markers (< 1 Mb apart) were detected in three of these fine-mapped loci in two QTL *(Growth4 on GGA3* and *Growth9 on GGA7)* in the backward elimination analysis. Below, we describe the dissection of these associations in more detail.

A major QTL, *Growth1*, was detected on GGA1 in the F_2_ intercross (Jacobsson et al. 2005; Wahlberg et al. 2009). The QTL-scan in the F_15_ generation of the AIL suggests that two loci, tagged by the markers *rs14916997 (p* = 2.1 ×10^-6^; likelihood ratio test) at 168.7 Mb and *rs14924102 (p* = 9.5 χ 10^-5^; likelihood ratio test) at 175.6 Mb, contribute to the F_2_ QTL (Figure 2A). The association to the marker *rs14916997* at 168.7 Mb was also detected in the multi-locus bootstrap-based backward elimination analysis *(p* = 4.2 × 10^-7^; likelihood ratio test; 5% FDR level). Although the multi-locus backward-elimination analysis also detected a second marker in this QTL, it was not the same one as in the QTL-scan, but to marker *rs316102705* at 171.5 Mb (*p* = 1.8 × 10^-1^; likelihood ratio test; 20% FDR; Table 1). A possible explanation for why an association is present in the QTL-scan, but not in the bootstrap-based backward-elimination analysis, is that its effects are dependent on the presence of alleles at other loci. This epistasis hypothesis is further explored in a separate section below.

The *Growth6* QTL extended across a long region of GGA4 in the F_2_ analysis (Jacobsson et al. 2005; Wahlberg et al. 2009) (Figure 2C). In the AIL F_15_, the QTL-scan (Figure 2C) and the multi-locus bootstrap-based backward-elimination analysis (Table 1) identify the same two regions to be associated with 56-day body-weight in the AIL F_15_ generation. In generation 40, the first region is centered around a marker at the proximal end of the QTL *(rs14417942* at 1.5 Mb; *p* = 6.5 × 10^-2^; likelihood ratio test) that is fixed for one allele in the HWS, and nearly fixed for the opposite allele in LWS (Table 1). At generation 50, this major LWS allele was fixed. The second association was to a marker located at the distal end of the QTL *(rs314557363* at 11.7 Mb; p = 7.1 × 10^-2^; likelihood ratio test) that was fixed for alternative alleles in the HWS and LWS lines at generation 40 (Table 1). The F_2_ *Growth6* QTL extends across these two fine-mapped loci in the F_2_, with the main peak in the middle (Figure 2C). The fine-mapping in the F_15_ therefore indicates that *Growth6* was a ghost-QTL (Knott and Haley 1992) in the F_2_ generation caused by the extended LD in this population.

The most significant QTL in the F_2_, *Growth9,* is located on GGA7 (Figure 2D). In the AIL F_15_, the QTL-scan and bootstrap-based backward-elimination analyses identified, 8 Mb apart, the same two independent loci in this QTL. The backward-elimination analysis detected associations to two markers in each of these loci (Figure 2D; Table 1). The proximal fine-mapped locus (16 Mb GGA7; galGal4) is located on the border of *Growth9,* whereas the distal locus (24 Mb GGA7; galGal4) is located in the middle of the QTL. The first peak at 16 Mb included associations to *rs14610961* (15.6 Mb; *p* = 4.0 × 10^-4^; likelihood ratio test) and *rs15853763* (16.2 Mb; *p* = 6.2 × 10^-3^; likelihood ratio test). In generation 40, both of these markers were segregating in HWS (Table 1). The second peak at 24 Mb included associations to *rs316269048* (23.9 Mb; *p* = 5.4 × 10^-3^; likelihood ratio test) and *rs315233740* (24.0 Mb; *p* = 2.1 × 10^-4^; likelihood ratio test). The first of these markers segregated in HWS in generation 40 (Table 1). A possible explanation for why associations are detected to multiple physically close markers (<1Mb apart) is that together they tag haplotypes with different effects that segregate at these loci. We describe how this hypothesis was tested for the loci where such associations were detected in a separate section below.

The *Growth12* QTL was mapped to GGA20 in the F_2_ population (Jacobsson et al. 2005; Wahlberg et al. 2009). The QTL-scan in the AIL F_15_ generation detected associations to markers in two linked loci on in this locus: *rs14278283* (10.5 Mb; *p* = 1 × 10^-3^; likelihood ratio test) and *rs14280503* (12.8 Mb; *p* = 3 × 10^-3^; likelihood ratio test) (Figure 2E). Only the association to *rs14280503* (5% FDR; Table 1) was significant in the bootstrap-based, backward-elimination analysis. Epistasis is a possible explanation for this finding and this was (for the locus detected in *Growth1*) tested in a separate section below.

### Epistatic interactions between markers associated with 56-day body-weight

As mentioned above, epistasis is a possible explanation for why some associations detected in the QTL-scan are not detected in the bootstrap-based backward-elimination analysis. This could be a result if the marginal effects of epistatic loci are dependent both on the allele-frequency at the locus itself and on the allele-frequency at other interacting loci elsewhere in the genome. As a consequence of this, the marginal additive effects of epistatic loci will be more sensitive to the bootstrapping procedure and hence potentially selected in fewer of the resampled populations. We therefore tested for epistatic interactions between the loci that were significant in the single marker association scan, but not in the bootstrap-based backward-elimination analysis. The pair *rs14924102* (in *Growth1*) and *rs14278283 (in Growth12)* displayed a significant epistatic interaction (p = 8.9 × 10^-3^). Their additive effects were also significant when fitted together with the 24 markers retained in the backward-elimination with bootstrapping analysis *(p* = 3 × 10^-3^ for *rs14924102* and *p* = 2.2 × 10^-4^ for *rs14278283;* Table 1, Table S1). The allele-frequencies at the marker *rs14924102* had, however, drifted from 0.5 during the breeding of the AIL *(p* = 0.13 in AIL F_15_). This resulted in a small number of observations in the two-locus genotype-classes including the minor-allele (A) at this locus (n = [3,11,8] for the [AAAA, AATA, AATT] genotypes). Hence, although there was a significant statistical interaction between these two loci, this result should be interpreted with caution: the epistatic interaction is due to the few individuals with the minor-allele homozygote genotype at this locus having opposite effects to those in the more frequent genotypes at this locus (AAAA and AATT; Supplemental Figure S1).

### Explorations of three fine-mapped loci in two QTL that segregate for multiple alleles

The backward-elimination analysis detected associations to pairs of physically close markers (<1Mb apart) at three loci in two QTL *(Growth4* and *Growth9).* In *Growth4* on GGA3 the associated markers are located 0.2 Mb apart and in *Growth9* on GGA7 they are located 0.6 and 0.1 Mb apart, respectively (Table 1). Because at least one marker in each pair segregate within one of the founder lines (Table 1), in practice, their two-locus genotypes act as multiallelic markers in the AIL. This allows tagging the haplotypes that segregate within the founder-lines to these markers and estimating their individual haplotype effects.

The proximal locus in the *Growth9* QTL on GGA7 was fine-mapped to two SNP markers 593 kb apart: *rs14610961* (15.6 Mb; *p* = 4.0 × 10^-4^; Table 1) and *rs15853763* (16.3 Mb; *p* = 6.2 × 10^-3^; Table 1). Both markers segregated for two alleles in HWS. Identified were three haplotypes across these two loci in the F_15_ population that segregated at frequencies > 0.1 *freq_AA_* = 0.35, *freq_GA_* = 0.10, and *freq_GG_* = 0.55). In the founders of this pedigree, that were from generation 40 of the selected lines, the LWS was fixed for GG, whereas HWS segregates for AA, AG and GG at frequencies *freq_AA_* = 0.65, *freq_AG_* = 0.175 and *freq_GG_* = 0.175, respectively (Supplemental Figure S2; Supplemental Table S2). Fitting a two-locus haplotype-ANOVA to these markers, while correcting for the effects of the other 22 selected markers (Table 1; Supplemental Table S1), did not reveal any significant associations between these multi-marker genotypes and 56-day body weight in the F_15_ (*p* = 0.08). That these marker haplotypes do, however, not fully capture the allelic complexity in the HWS and LWS lineages at generation 40 and 53 (Supplemental Figure S2), suggests that the region should be evaluated further using more informative markers to understand which haplotypes contribute to the multi-marker association at this locus.

The distal locus in the *Growth9* QTL was fine-mapped to two SNPs located 102 kb apart on GGA7: *rs316269048* (23.9 Mb, *p* = 4.0 × 10^-4^; Table 1) and *rs315233740* (24 Mb, *p* = 6.2 × 10^-3^; Table 1). The SNP *rs316269048* segregates for two alleles in HWS. The other SNP *rs315233740* is fixed for alternative alleles between HWS and LWS. Three haplotypes segregate at frequencies > 0.1 in the AIL F_15_ population *(freq_AA_* = 0.40, *freq_CA_* = 0.10, and *freqcT* = 0.48). In the founder lines at generation 40, the LWS is fixed for the CT haplotype, whereas HWS segregates for CT and AA at *freq_CT_* = 0.125 and *freq_AA_* = 0.875, respectively (Supplemental Figure S3; Supplemental Table S2). The CA haplotype in the F_15_, that was not observed in the founders, is likely a recombinant between the CT and AA haplotypes. The two-locus haplotype-ANOVA detected a significant association to 56-day weight in the F_15_ AIL population at this locus *(p* = 0.008). However, as the CT haplotype tags three different haplotypes that segregate at this locus in LWS at generation 40 (Supplemental Figure S3; Supplemental Table S2), and few individuals in the F_15_ carried each haplotype, the testing of individual haplotype effects were underpowered. Additional data are needed to explore in further detail the allelic heterogeneity at this locus.

The *Growth4* QTL on GGA3 was fine-mapped to two SNP markers 260 kb apart: *rs15321683* (33.6 Mb, *p* = 1.7 × 10^-5^; Table 1) and *rs14339371* (33.8 Mb, *p* = 1.9 × 10^-2^; Table 1, Figure 2B). The marker *rs15321683* was fixed for alternative alleles in HWS and LWS at generation 40. The marker *rs14339371* segregated for two alleles in LWS. Three haplotypes segregated at this locus in the F_15_ generation at frequencies > 0.1 *(freq_AG_* = 0.59, *freq_GA_* = 0.29, and *freq_GG_* = 0.12). The AG haplotype was almost fixed in HWS (p = 0.98), whereas GA and GG segregated in LWS at *(freqGA* = 0.61 and *freq_GG_* = 0.29, respectively; Table 3) at generation 40. The two-locus haplotype ANOVA detected a significant association to 56-day weight in the F_15_ AIL population in *Growth4 (p* = 0.001). The individual haplotype frequencies were sufficiently high in *Growth4* to allow an association analysis to the individual haplotypes. This analysis revealed that the segregating haplotypes have different effects on 56-day body-weight in the AIL F_15_ generation (Table 3). The major LWS haplotype at generation 40 (GA) decreased body-weight by 14.8g *(p* = 0.039; Table 3; Figure 3A) compared to the nearly fixed HWS haplotype (AG). The minor LWS haplotype (GG) decreased 56-day weight even more (-44.3g; *p* = 0.0003; Table 3; Figure 3A).

**Table 3.**
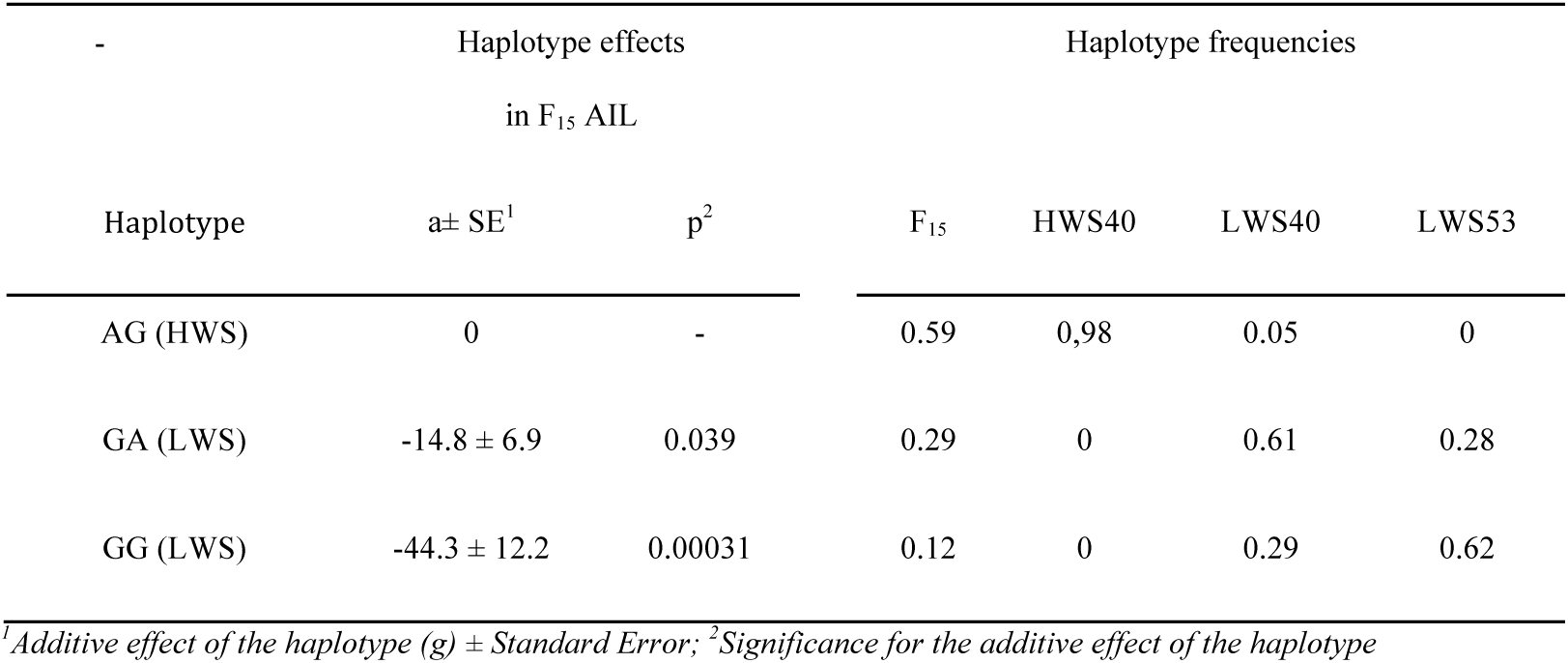
Estimated effects and allele-frequencies for the haplotypes segregating in *Growth4* on GGA3

We continued to explore the high-density SNP haplotypes in the HWS and LWS lineages across the region of *Growth4* that contained the two significantly associated linked SNP markers (GGA4: 32.1–35.9 Mb) in detail. The major two-locus HWS haplotype (AG) and the minor LWS haplotype (GG) at generation 40 each tag well-defined haplotypes across the region (Figure 3B; Figure S4). The major two-marker LWS-haplotype (GA) at generation 40, however, tags two different haplotypes. During selection for decreased body-weight in LWS from generation 40 to 53, the frequency of the haplotype with the lowest effect on 56-day weight (GG) increased (Figure 3B, Supplemental Figure S4). By generation 53, it has become the major haplotype in the LWS line (Table 2; Supplemental Figure 3B). This suggests an ongoing selection at this QTL at generation 40, leading to fixation of the haplotype with the least effect on 56-day body-weight by generation 53.

**Figure 3.**
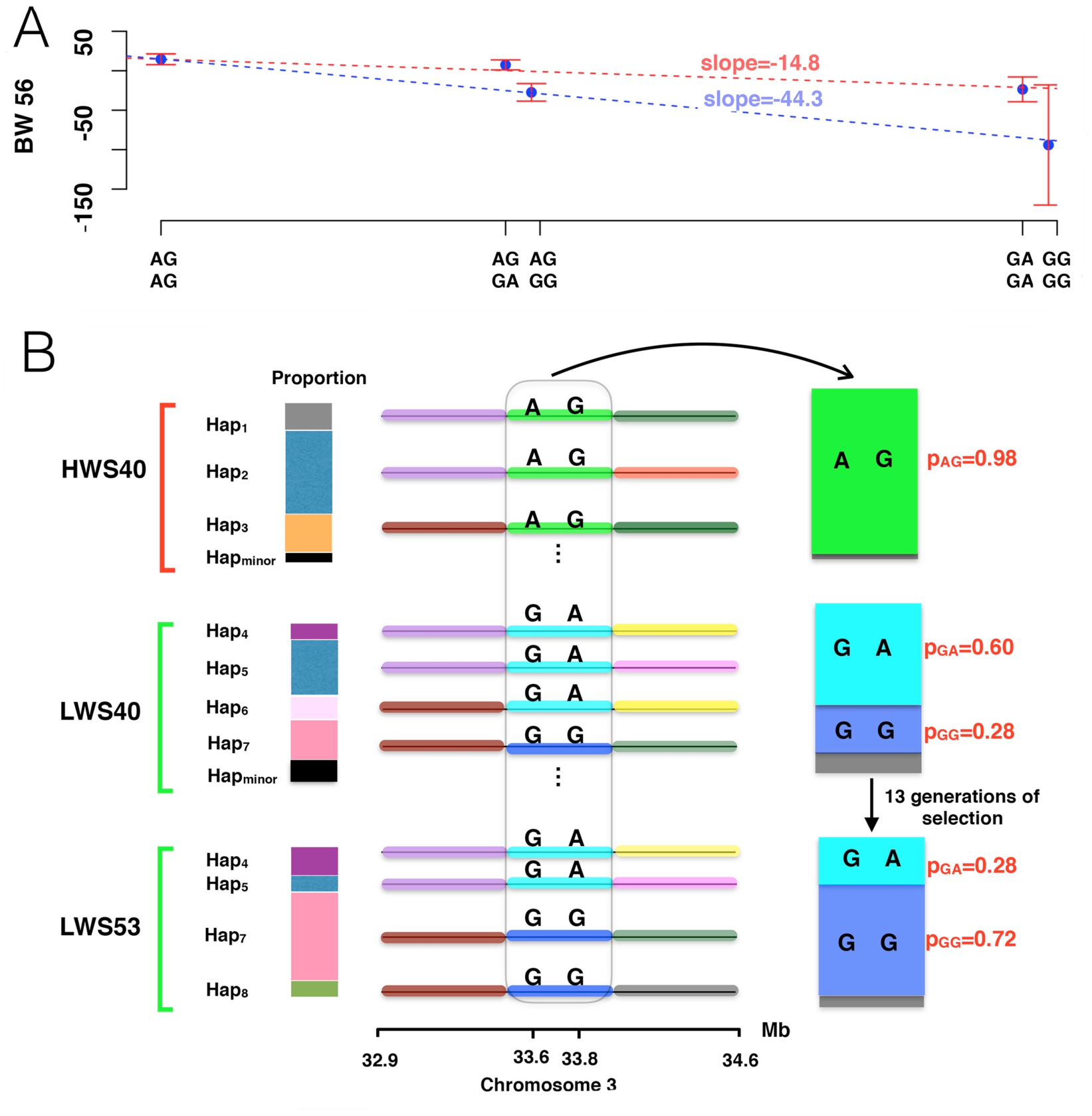
Schematic illustration of the haplotype-structure around markers rs15321683 (33.6 Mb; galGal4) and rs14339371 (33.8 Mb; galGal4) on GGA3, and how the founder-haplotypes that they tag contribute to 56-day body-weight in the AIL F_15_ generation. **(A)** At this locus, the individuals segregate for three major haplotypes that have distinct effects on 56-day body weight in the AIL F_15_ generation. The figure illustrates the regressions obtained for individuals carrying different haplotypes in the AIL F_15_ generation (the error-bars indicate the standard-errors of the estimates). **(B)** A schematic illustration of the major haplotypes segregating in the region from 32.9-34.6 Mb on GGA3 (a complete visualization of the haplotype structure based on all SNPs genotyped on the 60K SNP chip is provided in Supplemental Figure S4). The HWS line is fixed for a weight-increasing haplotype AG, while the LWS line segregates for two weight-decreasing haplotypes (GA, GG) in the **QTL** *Growth4* on GGA3 at generation 40. The haplotype in the LWS lineage with the strongest body weight decreasing effect tags a single haplotype that increases in frequency during selection from generation 40 to 53.

### Variance explained by the fine-mapped loci

We estimated how much of the additive genetic variance for 56-day body weight in the AIL F_15_ population that could be explained by the associated markers in the fine-mapped QTL (Table 1) and those identified in selective-sweep regions elsewhere in the genome (Sheng et al. 2015) (Supplemental Table S1). Together 24 markers, representing 20 loci, retained in the backward-elimination analysis (20 % FDR) explained 27.8% of the residual phenotypic variance, corresponding to 60.5% of the additive genetic variance in the population.

## Discussion

In this study, we have continued to dissect the genetic architecture underlying the highly polygenic trait, 56-day body-weight, in the Virginia chicken population that has been subjected to long-term, divergent single-trait selection. With a focus on the F_15_ generation of an AIL between the lines, we fine-mapped and explored the genetics of nine major QTL that were mapped in the F_2_ (Jacobsson et al. 2005; Wahlberg et al. 2009) and replicated in the F_2_ to F_8_ generations (Besnier et al. 2011; Brandt et al. 2016) The increased resolution in the F_15_ population allowed us to fine-map several of the major QTL to contributions by linked loci, and sometimes multiple haplotypes within and epistatic interactions across these loci. These findings further emphasize the important contribution by many loci with small additive effects to adaptation (Jacobsson et al. 2005; Lango Allen et al. 2010; Wood et al. 2014), where some of them are tightly linked (Iraqi *et al*. 2000). We also found that one of the original QTL detected in the F_2_ population (*Growth6* on GGA4) was likely a ghost-QTL (Haley and Knott 1992).

Our backward-elimination based mapping approach revealed associations to markers that were physically close but segregated for multiple alleles in at least one of the founder lines. By utilizing haplotype information on the founders, we confirmed that these markers tag segregating haplotypes with distinct effects on the traits and provide insights to the potential importance of allelic heterogeneity (Forsberg et al. 2015; McClellan and King 2010; Yano et al. 2016) for the response to long-term selection in the Virginia lines. Because the current population is rather small, the haplotype-frequencies have drifted away from 0.5 during the breeding of the deep intercross population (minor allele frequency ranging from 0.03 to 0.5, with median of 0.39). In addition, the markers in this study were not initially selected to tag the segregating haplotypes, resulting in low statistical power to test or estimate the effects of all individual haplotypes. Further studies to explore these multi-allelic loci with more informative SNP markers would help disentangle the likely important contributions by allelic heterogeneity in the fine-mapped loci to the long-term selection response.

Previous studies of the Virginia body-weight selected populations revealed and replicated extensive epistatic interactions among QTL alleles (Carlborg et al. 2006; Le Rouzic et al. 2008). Unfortunately, the allele-frequencies were too imbalanced and the number of individuals too small in this F_15_ dataset to provide sufficient statistical power for an extensive exploration of epistasis. Therefore, we limited testing of epistasis to explore the possibility that two of the loci found to be sensitive to the genetic background in the bootstrap analysis are involved in an interaction.

The 24 markers in the fine-mapped QTL and selective sweeps outside QTL regions explained 27.8 % of the residual phenotypic variance or 60.5% of the additive genetic variance in the F_15_ population. This is considerably more than the 13% explained by the single significant and 12 suggestive QTL in the original F_2_ line-cross analysis (Jacobsson et al. 2005). In the F_2_-F8 generations of the AIL, the five most significant QTL explained 10% of the residual phenotypic variance (Besnier et al. 2011). Our dissection of this adaptive trait in this long-term selection experiment has thus revealed loci that explain nearly two thirds of the additive genetic variance in this population. Although a large fraction of this variance is contributed by loci outside the QTL (Sheng et al. 2015), the linked and segregating loci revealed in the fine-mapping of the QTL suggest that much genetic variance has been released due to recombination break-up of unfavorable linkages in the selected lines during the breeding of the AIL. One example of this is the mapping of transgressive alleles in the F_15_ (e.g. between *rs316102705* and *rs14916997* in *Growth1* on GGA1; Table 2). These unfavorable linkages will decrease the genetic variance explained by the QTL in the F_2_ where the LD extends over both these loci.

For the QTL that were fine-mapped to a single bi-allelic locus, there was an overall agreement between the additive effects estimated in the F_2_ and F_15_ generations (Table 2). This was consistent with expectations that there should be less bias in the estimates of QTL effects when sufficiently large QTL mapping populations are used (Beavis 1998). For the QTL with more complex genetic architectures, including those with linked loci and multiple alleles, the deviation was larger, as the allele substitution effect at a single marker is not sufficient for representing such effects (Table 2). The degree to which these estimates differ will depend, for example, on the linkage phase of the linked loci and the frequencies of the respective alleles at the multi-allelic locus. Our results show that overall the marginal effects estimated for the F_2_ QTL agree with the joint estimated effects for the fine-mapped loci and alleles in the F_15_ generation. Further work is needed to explore whether the remaining deviations are due to still unresolved complexities for these loci; for example incomplete tagging of segregating alleles in the founders, multi-allelic genetic architecture, and/or epistatic interactions with other loci.

The ultimate aim for studies designed to dissect the genetic architecture of a complex trait is to find the causal genes and mutations underlying the trait. Although the use of a deep intercross population has allowed us to fine-map the major QTL segregating in this population to 1-2 Mb resolution, the limitation of long LD blocks in linkage mapping analysis still limited our ability to achieve this aim. Additional research is necessary to identify haplotypes segregating within and across the selected lines and AIL, to reduce the number of candidate genes in the 24 identified loci. Regardless, this provided valuable genome-wide insights to the variety of genetic mechanisms that contribute to the genetic variance for a single trait for which dramatic adaptations have emerged during long-term directional selection.

In conclusion, this study illustrates that the strong response to long-term divergent selection in the Virginia body weight lines rely on a highly polygenic and complex genetic architecture. Most of the original major QTL in this population fine-map to contributions by linked, and sometimes multi-allelic or epistatic loci. By continuing the in-depth characterizations of the highly polygenic genetic architecture of this adaptive trait, we provide insights for future research aiming to understand the variety of genetic mechanisms that together are likely to contribute to adaptation and evolution. It demonstrates that, even in relatively small, closed populations, selection response is likely to involve a wide variety of the complex genetic mechanisms that together contribute to quantitative trait variation and adaptation.

## Materials and Methods

### Ethics statement

All procedures involving animals used in this experiment were carried out in accordance with the Virginia Tech Animal Care and Use Committee protocols.

### Animals, phenotyping and DNA extraction

The chickens used in this study were from the F_15_ generation of an Advanced Intercross Line bred from the Virginia high (HWS) and low (LWS) body weight selected lines. The HWS and LWS lines were founded in 1957 from a common base population obtained by crossing seven partially inbred lines of White Plymouth Rock chickens. They have since then been subjected to bi-directional selection for high or low 56-day body weight, respectively. For further details on the Virginia lines, see Dunnington et al (Dunnington and Siegel 1996; Dunnington et al. 2013). An Advanced Intercross Line (AIL) was founded from generation 40 of the HWS and LWS, where the sex-average 56-day body-weights were 1,412 g (SE: ± 36 g) and 170 g (SE: ± 5 g), respectively (Jacobsson et al. 2005). This AIL has previously been used in the fine-mapping of QTL using the F_2_-F_8_ AIL generations (Besnier et al. 2011; Brandt et al. 2016; Pettersson et al. 2011). In the AIL F_15_ generation, 907 individuals were hatched and 56-day body weights and genotypes measured on 825 (Genotype and phenotype data are available as supplemental data 1 and 2). These birds were used in an association analyses to evaluate contributions by genome-wide selective sweeps to 56-day body-weight the F_15_ generation (Sheng et al. 2015). This study is based on the same individual F_15_ individuals as in Sheng et al (Sheng et al. 2015).

### Marker selection and genotyping

In total, 434 SNP markers, genotyped using a GoldenGate genotyping assay (Illumina Inc; performed at the SciLifeLab SNP&SEQ Technology Platform at Uppsala University), were included in this study. Of these, 252 markers were genotyped and used in an earlier study of the AIL F_15_ generation (Sheng et al. 2015) to target 99 regions in the genome where the HWS and LWS lines were fixed for alternative alleles after 50 generations of selection. These 99 regions cover about 138 Mb of the 1.2 Gb in the chicken reference genome (galGal4). For this study, we genotyped an additional 182 SNP markers selected to provide dense marker coverage across nine QTL known to affect body weight in the F_2_ (Carlborg et al. 2006; Jacobsson et al. 2005; Wahlberg et al. 2009) and F_2_-F_8_ intercross populations (Besnier et al. 2011; Brandt et al. 2016; Pettersson et al. 2011). Together the 434 markers covered approximately 300 Mb of the chicken genome (galGal4), of which 170 Mb represent targeted QTL regions (in total 216 markers), and 130 Mb (218 markers) selective-sweep regions (Sheng et al. 2015). The Nov 2011 (galGal4) chicken genome assembly was used for comparing physical locations of mapping results from the F_2_ (Wahlberg et al. 2009) and the F_15_ intercross generations.

### Coding of SNP marker alleles in the AIL F_15_ generation

The GoldenGate assay reports SNP marker alleles on a [A, T, C, G] basis. Before the statistical analysis, we re-coded the marker genotypes in individuals as [-1, 0, 1]. The coding -1/1 was used to represent homozygotes for the most common alleles in the LWS/HWS founders, respectively. The code 0 was used to represent heterozygotes. The LWS/HWS origin was determined by comparing the F_15_ genotypes to those of the HWS/LWS F_0_ founders. By using this coding, positive estimates of additive effects in the association analysis indicate that predominant allele in HWS increased weight, and negative additive **effects that the predominant allele in LWS increased weight.**

### Multi-locus association analysis using a backward-elimination strategy with bootstrapping to correct for population structure

The statistical analyses were designed to simultaneously fine-map nine QTL regions contributing to 56-day body-weight in the Virginia body weight lines, while also accounting for the effects of other regions across the genome. This was to appropriately account for the highly polygenic genetic architecture of body-weight in this population (Besnier et al. 2011; Jacobsson et al. 2005; Johansson et al. 2010; Pettersson et al. 2013; Sheng et al. 2015), where many of the genotyped markers both in and outside the QTL regions should contribute to the trait. As all individuals in the AIL were progeny of dams of the same age, hatched on the same date, and reared separate from their parents, the environmental contributions to between-family means in the F_15_ population should be minimal. We therefore assume that a large portion of the differences in family means are due to the joint effects of the markers in the genotyped QTL and selective sweep regions, rather than non-genetic effects. Population structure might, however, still be of concern in a deep-intercross population (Peirce et al. 2008; Cheng et al. 2010). We therefore also validate our results using a bootstrap-based approach developed to account for the possible effects of population structure in general deep-intercross populations, including AILs (Valdar et al. 2009).

Not known in advance was how many genetic markers in the fine-mapped QTL contributed to 56-day body-weight in the AIL F_15_ generation. Based on previous findings (Besnier et al. 2011; Brandt et al. 2016), we expected that at least some QTL would contain multiple linked loci. The number of loci in the final model could therefore vary substantially, and to account for this we implement our analysis with an adaptive model selection criterion controlling the False Discovery Rate (FDR) (Abramovich et al. 2006; Gavrilov et al. 2009) developed for this purpose. Starting with the 434 genotyped SNP markers, we then selected a final set of markers using a two-step backward-elimination strategy applied on a standard linear model with 56-day body-weight as response variable. A 20 % FDR among the selected loci was used as the termination criterion.

In the first step, a linear model including the SNP markers to be tested, together with the fixed effects of sex of the individual and the genotypes of the 16 sweeps that were earlier found to be associated with 56-day weight in this population (Sheng et al. 2015). The 16 earlier mapped sweeps were included in the model to capture the major polygenic effect in this population. To avoid over parameterizing the linear model, the 434 markers were divided into eight pre-screening sets including approximately 50 markers each. These sets were selected to, as far as possible, include markers from the same chromosomes. A backward-elimination analysis with the adaptive FDR criterion was performed on each set separately. All markers that reached the 20% FDR in either of these analyses were kept for the further joint analyses described in the next section.

In the second step of the analysis, all markers selected in the pre-screening were analyzed jointly using the bootstrap based method of Valdar et al (Valdar et al. 2009). This analysis was used to identify the loci that contribute to 56-day body weight in this population and that was robust to the possible effects of population structure. Here, a RMIP (Resample Model Inclusion Probability) was calculated for each marker tested. A final model was selected where only markers with RMIP > 0.46, the threshold suggested for an AIL generation F_18_ (Valdar et al. 2009), was included. All of these analyses were implemented in custom scripts using the statistical software R (R Core Team 2015).

### Estimation of the additive effects and significances of the markers associated with 56-day body-weight in the F_15_ population

The general linear model used to estimate additive effects and significances of the loci associated with 56-day body-weight in this population can be formulated as

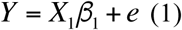

Here, *Y* is a column vector containing the 56-day body-weight of the 825 F_15_ individuals. *β_1_* is a vector with the estimate of the fixed effect of sex and the additive effect of all markers tested. *X_1_* is the design matrix including the coding for the sex of the birds and the genotypes of the markers coded as [-1,0,1]. *e* is the normally distributed residual. The markers included to control for the background genetic effects will vary depending on the analysis as described in the sections below.

### Analyses of markers selected in the backward-elimination analysis

There are few recombination events within each QTL in the AIL pedigree which limits the resolution in the fine-mapping analysis. In the backward-elimination analysis, we therefore consider markers located closer than 1 Mb from each other as representing the same fine-mapped locus. The effects of all markers in these loci were estimated using the model described above (model 1) by defining *X_1_* to include only the genotypes of the tested marker from that locus and the SNPs in the remaining loci. This meant that when estimating the effect of a maker located inside a fine-mapped locus tagged by more than 1 marker, the remaining markers inside this locus were excluded from the analysis. The significance for each marker was obtained using a likelihood ratio test comparing regression models (model 1) with and without the tested marker.

### Single marker association scans in the replicated QTL

To obtain statistical support profiles for the QTL with significant associations in the backward-elimination analysis, we performed single marker association scans across all markers in these QTL. The same regression model (model 1) was used, but *X*_1_ here include the genotype of the tested marker together with the sex effect and the additive effects of all SNPs selected in the backward-elimination analysis.

### Effects of the QTL detected in the F_2_ population

The additive effects and significances for QTL identified in the F_2_ population were extracted at the location of the marker closest to the reported QTL-peak from the results in (Wahlberg et al. 2009).

### Calculation of founder-line allele-frequencies for the markers associated with body-weight in the F_15_ population

Previously, we genotyped 20 individuals from each of the HWS and LWS lines from generation 40 using a 60k SNP chip (Jacobsson et al. 2005; Pettersson et al. 2013). Some of the SNPs evaluated here were included in that study and for those SNPs those data were used to calculate the allele-frequencies in the founder-lines. For the remaining SNPs, the allele-frequencies in the founder-lines were estimated using sequence data (∼30X coverage) from two pools including 29 HWS founders and 30 LWS founders, respectively (Supplemental data 3 and 4).

### Testing for pairwise epistatic interaction between loci

It is possible that association peaks that are significant in the scan across the QTL in the original data, but not in the bootstrap-based backward-elimination analysis, could be lost due to sensitivity to allelic background at other loci (i.e. epistasis). We therefore tested for epistasis between such peaks, using a likelihood ratio test, by evaluating the significance of the pairwise interactions in model 2:

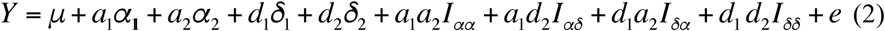

where *Y* is the residual from a regression model (model 1) fitting all significant markers selected in the bootstrap-based backward-elimination analysis together with the sex of the chickens. μ is the reference point, *α_1_* and *α_2_ (δ_1_* and *δ_2_)* are the additive allele-substitution effects (dominance deviations) at the two loci, *I_αα_, I_αδ_* and *I_δδ_* are the additive by additive, additive by dominance, dominance by dominance and dominance by dominance interaction effects, and *e* is the normally distributed residual. The *a\** and *d\** are the NOIA indicator regression variables for the two loci. These indicator regression variables are allele-frequency weighted codings of the genotypes at the evaluated loci and calculated as detailed in (Alvarez-Castro and Carlborg 2007). Using the orthogonal statistical NOIA parameterization of the model, we tested for epistasis by comparing the full model (model 2) to the reduced model without the interaction effects using a likelihood ratio test. All models were fitted using the *lm* function in R with design matrices created by the *multilinearRegression* function in *noia* R-package (Alvarez-Castro and Carlborg 2007; Le Rouzic and Álvarez-Castro 2008). P values were calculated using the *lrtest* function in the R-package *lmtest* (Zeileis and Hothorn 2002).

## Analyses of segregating haplotypes

### Haplotype estimation in individuals from the founder-lines and AIL F_15_ generation

We inferred the haplotypes for n = 20 individuals from each of generations 40 (founders) and 53 from the HWS and LWS lines (in total 80 birds), and all 825 birds with phenotypes from the AIL F_15_ generation. This was done using the software *fastPHASE* (Scheet and Stephens 2006) with default parameters “fastPHASE –Z input-file”. For this, we used 60k SNP chip genotypes generated for the generation 40 and 53 HWS and LWS individuals in two earlier studies (Johansson et al 2010) and the new 434 SNP markers genotypes in the AIL F_15_ individuals. Some of the SNP markers genotyped in the AIL F_15_ were not included on the 60k SNP chip and could therefore not be unambiguously assigned to HWS or LWS haplotypes based on this haplotyping. Instead, they were assigned to a founder haplotype based on the agreement between their allele-frequencies, estimated from the pooled whole-genome resequencing data described above, and those of a nearby SNP from the 60k SNP-chip genotype data.

### Haplotype-based association analysis

A haplotype-based association analysis was performed in regions of the QTL where the bootstrap-based backward-elimination analysis, and the haplotype estimation, suggested that multiple alleles were segregating in one, or both, of the founder lines. These analyses used a one-way ANOVA (model 1) where *Y* is a column vector including the residuals from a regression model fitted on the phenotypes of all individuals with the genotypes of all markers selected in the backward-elimination analysis located outside of the evaluated locus and sex as fixed effects. *X* is the design matrix with one column for the reference-point and multiple columns, and each contain a haplo-genotype. *β* is a column vector with the estimates for the reference point and the deviations from the reference point for each haplo-genotype, and *e* is the normally distributed residual.

### Estimation of genetic effects of individual haplotype effects

For loci where the haplotype-based association analysis indicated that multiple haplotypes were segregating in one, or both, of the founder lines, also performed was an analysis to disentangle the effects of the individual haplotypes. For a locus with multiple haplotypes (denoted here as A, *B, C….* N), we first identified the common haplotypes (p > 0.1) and only included them in the analysis. Second, a strata was formed that only contained homozygous and heterozygous individuals for particular haplotype combinations (e.g. *AA, AB, BB).* Third, using these strata we estimated the effects of the individual haplotypes, in turn (e.g. the effects of *A* and *B* were estimated in the strata containing individuals with the haplotype combinations *AA, AB* and *BB*) using a regression model (model 1) parameterized as follows. *Y* here contains the residual for the birds in an analyzed strata from a regression model where the genotype of all markers located outside of the tested locus, and sex of the birds, were fitted as fixed effects in a regression on the 56-day body-weight. *X* is here the design matrix where the haplo-genotype for individuals homozygous for the LWS derived haplotype, heterozygous for LWS/HWS derived haplotype and homozygous for the HWS derived haplotype are coded as [0,1,2], respectively. *e* is the normally distributed residual. This analysis was repeated for the all the major haplotypes for which sufficiently large numbers of individuals were genotyped and phenotyped (n > 50) in at least two of the three haplo-genotype classes in the AIL F_15_ generation. The significances for the associations obtained as the p-value from a Wald test calculated using the *lm* function in R.

## DataAccess

All raw data will be released upon publication of the final manuscript.

## Acknowledgements

We thank Leif Andersson for initiating the intercross experiment with PBS and sharing the QTL mapping data from the F_2_ intercross. Stefan Marklund, Andreas Lundberg and Mats Pettersson are acknowledged for their valuable contributions during selection of polymorphisms in the selective-sweep regions for genotyping, development of primers for these, and for interacting with the genotyping centre. We also thank Mats Pettersson for useful input on the analysis of the data. Genotyping was performed by the SNP&SEQ Technology Platform in Uppsala, which is part of Science for Life Laboratory at Uppsala University and is supported as a national infrastructure by the Swedish Research Council (VR-RFI). This study was funded by the Swedish Research Council (VR grant ID 2012-4634) and the Swedish Research Council for Environment, Agricultural Sciences and Spatial Planning (Formas grant IDs 2010-643 and 2013-450).

## Author contributions

ÖC and PBS initiated the study and designed the project; PBS developed the Virginia chicken lines; PBS designed, planned and bred the Virginia Advanced Intercross Line; PBS and CFH designed, planned, bred, bled, phenotyped, and extracted DNA from the F_15_ Virginia Advanced Intercross Line population; ZS performed the quality control of the genotype data; ÖC and LR designed the statistical analyses; ZS, YZ and ÖC contributed analysis scripts; ÖC, YZ and ZS performed the data analyses and summarized the results. ÖC and YZ wrote the manuscript. All authors read and approved the final manuscript.

## Disclosure declaration

The authors declare no competing interests.

### Legends for supplemental figures and tables

**Figure S1.**
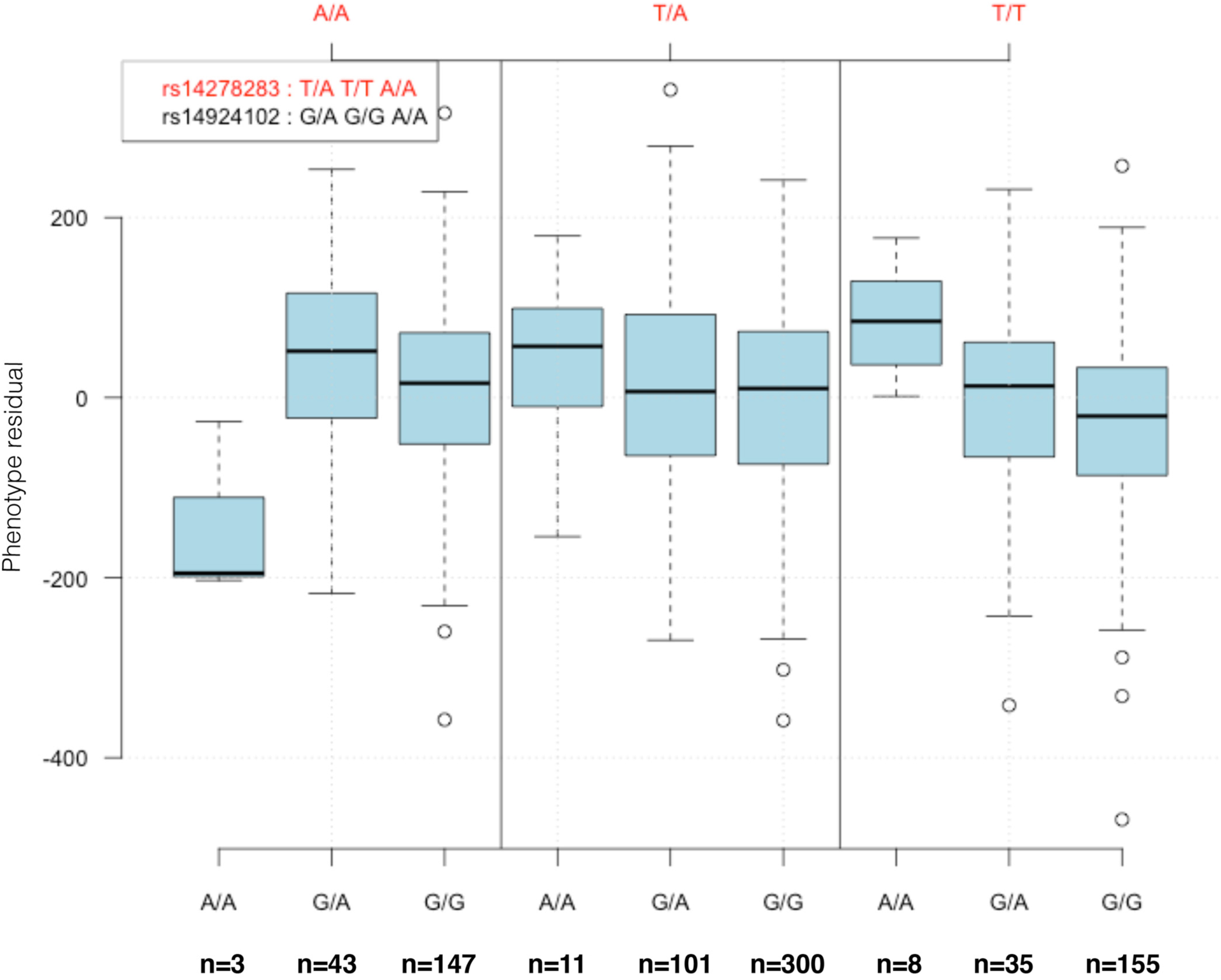
Two-locus genotype-phenotype plot. The Y-axis shows the residual phenotypes from a regression model including the 24 associated markers in the multi-locus association analysis. The two markers representing the loci, rs14924102 *(Growth1)* and rs14278283 (*Growth12*), were selected as they were the ones with the strongest marginal associations to 56-day body weight in the respective QTL.

**Figure S2.**
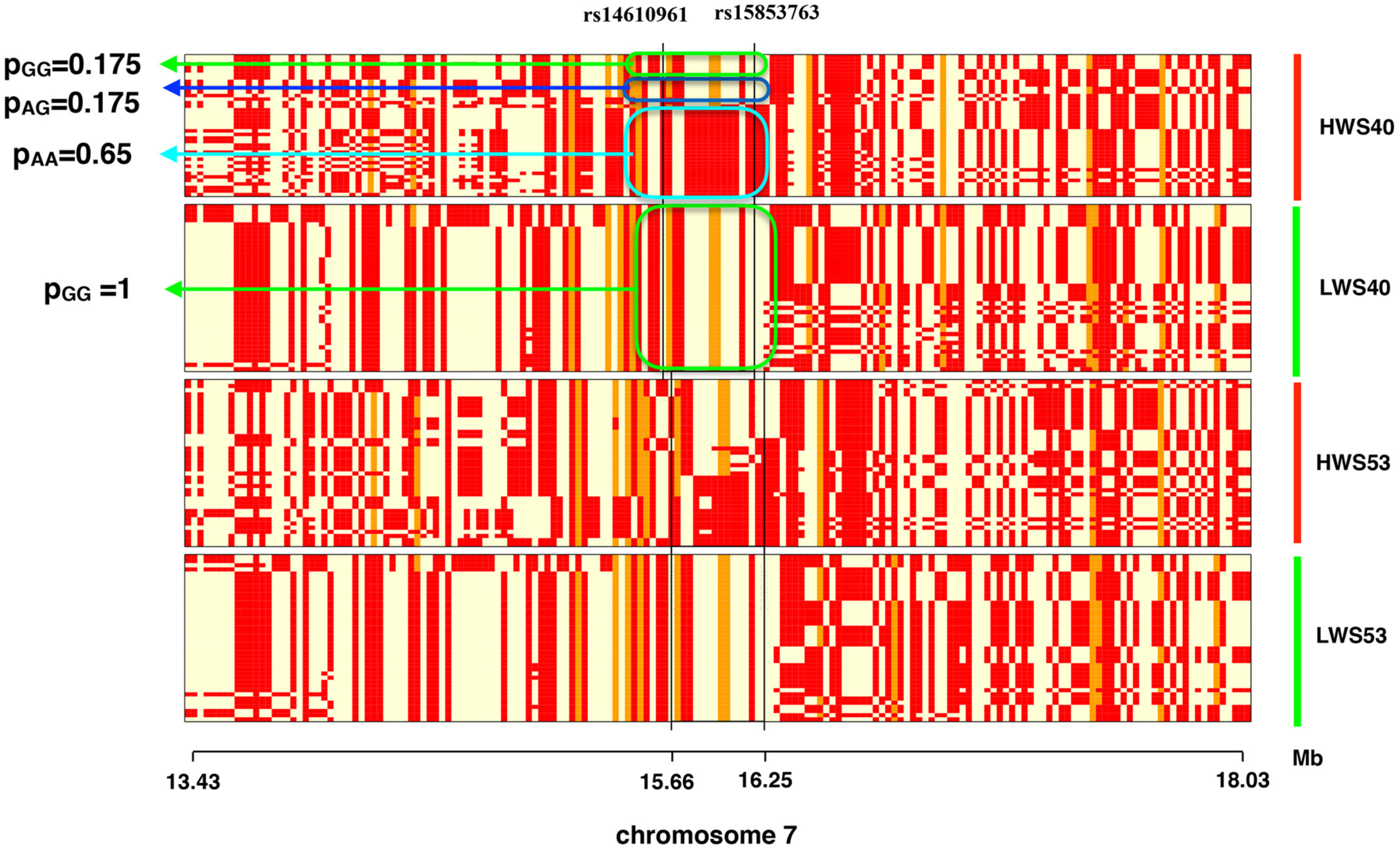
Visualization of the haplotypes inferred across the polymorphic 60k SNP-chip markers in the region from 13.4-18.0 Mb on GGA7 in the HWS and LWS lineages at generations 40 and 53. In every panel, the rows represent an individual chromosome and the columns the marker genotype at a SNP, where different genotypes are plotted in in different colours.

**Figure S3.**
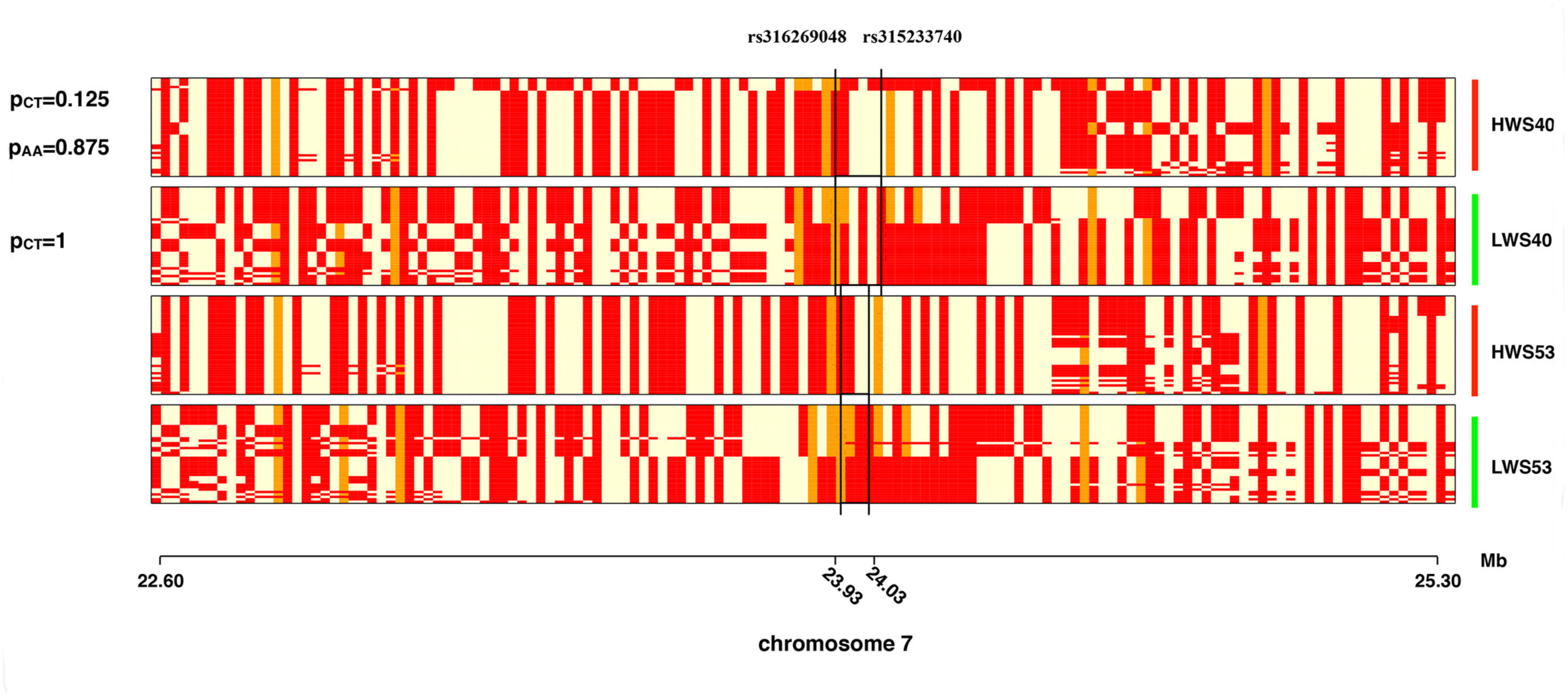
Visualization of the haplotypes inferred across the polymorphic 60k SNP-chip markers in the region from 22.6-25.3 Mb on GGA7 in the HWS and LWS lineages at generations 40 and 53. In every panel, the rows represent an individual chromosome and the columns the marker genotype at a SNP, where different genotypes are plotted in in different colours.

**Figure S4.**
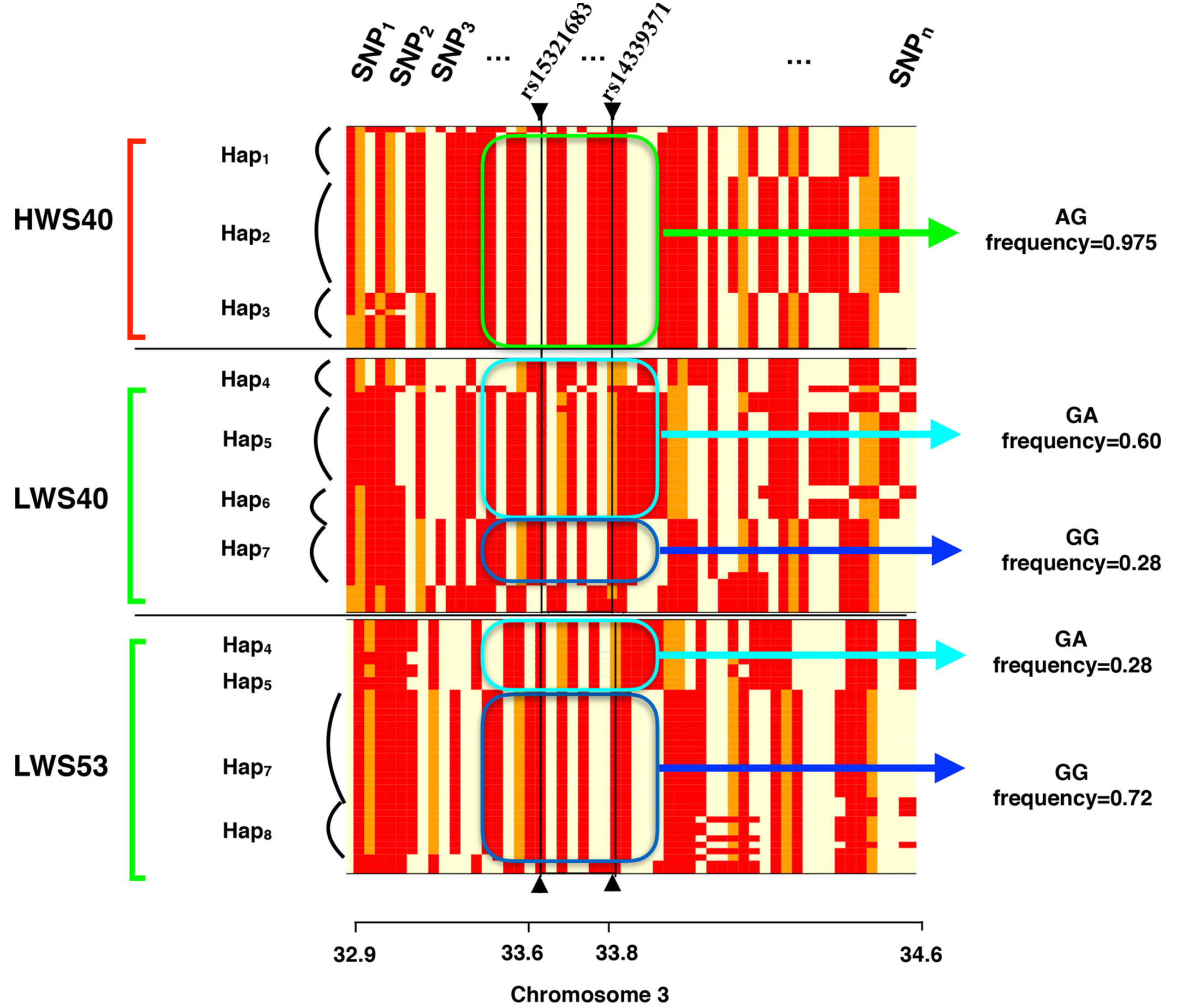
Visualization of the haplotypes inferred across the polymorphic 60k SNP-chip markers in the region from 32.9-34.6 Mb on GGA3 in the HWS and LWS lineages at generations 40 and 53. In every panel, the rows represent an individual chromosome and the columns the marker genotype at a SNP, where different genotypes are plotted in in different colours.

**Table S1.** The 11 SNP markers that are associated with 56-day body-weight, and located outside the nine fine-mapped QTL regions, in generation F_15_ of the Advanced Intercross Line between founders from generation 40 of the High-(HWS] and Low (LWS] body-weight selected Virginia chicken lines.

| GGA<sup>a</sup> | Pos<sup>b</sup> | Marker | Freq<br>HWS/LWS40<sup>c</sup> | a ± SE<sup>d</sup> | P value<sup>e</sup> | RMIP<br>5/20%<sup>f</sup> | Sign<sup>g</sup> |
| --- | --- | --- | --- | --- | --- | --- | --- |
| 1 | 33147325 | rs13847968 | 0.68/0* | -6.0±6.2 | 3.0 x 10<sup>-1</sup> | 0.33/0.46 | 0.20 |
| 1 | 87189315 | rs13899455 | 1/0.08 | -15.8±5.7 | 3.3e-03 | 0.53/0.64 | 0.05 |
| 1 | 132884851 | rs13942473 | 0.95/0 | 14.3±7.0 | 2.6e-02 | 0.57/0.70 | 0.05 |
| 2 | 148418221 | rs15158686 | 1/0 | 24.0±5.8 | 4.3e-06 | 0.66/0.76 | 0.05 |
| 4 | 69059781 | rs314510951 | 1/0 | 9.6±5.4 | 5.1e-02 | 0.37/0.53 | 0.20 |
| 4 | 82310849 | rs14498744 | 1/0 | -21.2±5.9 | 7.7e-05 | 0.74/0.85 | 0.05 |
| 6 | 3292946 | rs13562096 | 1/0 | 12.7±5.9 | 3.4e-05 | 0.60/0.70 | 0.05 |
| 6 | 3417847 | rs15758084 | 1/0 | 1.9±5.9 | 5.0e-04 | 0.60/0.70 | 0.05 |
| 10 | 8504427 | rs312978808 | 1/0 | 10.6±5.6 | 2.9e-02 | 0.40/0.58 | 0.20 |
| 13 | 5988992 | rs13726878 | 0.95/0 | 14.7±5.9 | 7.4e-03 | 0.46/0.62 | 0.20 |
| 23 | 4599103 | rs15205573 | 1/0.09 | 15.1±6.2 | 7.2e-03 | 0.58/0.76 | 0.05 |

*^a^GGA:* Gallus Gallus Autosome, ^b^Pos:November 2011 (galGal4) assembly; ^c^Freq HWS/LWS: Estimated allele frequency in the HWS/LWS founders using individual SNP chip genotypes or, if this was not available, pooled sequencing data (labeled with *); ^d^Additive genetic effect ± *Standard Error; ^e^Significance for additive genetic effect in model including all loci significant at 20 % FDR;^f^Resample Model Inclusion Probability at 5/20 % FDR threshold; ^g^FDR threshold at which the marker was selected with RMIP >0.46[11].*

**Table S2.** Inferred haplotypes and their allele-frequencies in the two 56-day body weight associated regions in the QTL *Growth9* on GGA7.

| QTL region | Haplotypes | Haplotype frequencies |  |  |
| --- | --- | --- | --- | --- |
|  |  | F15 | HWS40 | LWS40 |
| Growth 9.1 | AA (HWS1) | 0,35 | 0,65 | 0 |
|  | GA (HWS2) | 0,09 | 0 | 0 |
|  | AG (HWS3) | 0 | 0,175 | 0 |
|  | GG (LWS1) | 0,55 | 0,175 | 1 |
| Growth 9.2 | AA (HWS1) | 0,4 | 0,875 | 0 |
|  | CA (HWS2) | 0,1 | 0,125 | 0 |
|  | CT (LWS1) | 0,48 | 0 | 1 |

Haplotypes and their corresponding allele-frequencies for two regions in Growth9

